# ViralLink: An integrated workflow to investigate the effect of SARS-CoV-2 on intracellular signalling and regulatory pathways

**DOI:** 10.1101/2020.06.23.167254

**Authors:** Agatha Treveil, Balazs Bohar, Padhmanand Sudhakar, Lejla Gul, Luca Csabai, Marton Olbei, Martina Poletti, Matthew Madgwick, Tahila Andrighetti, Isabelle Hautefort, Dezso Modos, Tamas Korcsmaros

## Abstract

The SARS-CoV-2 pandemic of 2020 has mobilised scientists around the globe to research all aspects of the coronavirus virus and its infection. For fruitful and rapid investigation of viral pathomechanisms, a collaborative and interdisciplinary approach is required. Therefore, we have developed ViralLink: a systems biology workflow which reconstructs and analyses networks representing the effect of viruses on intracellular signalling. These networks trace the flow of signal from intracellular viral proteins through their human binding proteins and downstream signalling pathways, ending with transcription factors regulating genes differentially expressed upon viral exposure. In this way, the workflow provides a mechanistic insight from previously identified knowledge of virally infected cells. By default, the workflow is set up to analyse the intracellular effects of SARS-CoV-2, requiring only transcriptomics counts data as input from the user: thus, encouraging and enabling rapid multidisciplinary research. However, the wide-ranging applicability and modularity of the workflow facilitates customisation of viral context, *a priori* interactions and analysis methods. Through a case study of SARS-CoV-2 infected bronchial/tracheal epithelial cells, we evidence the functionality of the workflow and its ability to identify key pathways and proteins in the cellular response to infection. The application of ViralLink to different viral infections in a cell-type specific manner using different available transcriptomics datasets will uncover key mechanisms in viral pathogenesis. The workflow is available on GitHub (https://github.com/korcsmarosgroup/ViralLink) in an easily accessible Python wrapper script, or as customisable modular R and Python scripts.

**Author summary:** Collaborative and multidisciplinary science provides increased value for experimental datasets and speeds the process of discovery. Such ways of working are especially important at present due to the urgency of the SARS-CoV-2 pandemic. Here, we present a systems biology workflow which models the effect of viral proteins on the infected host cell, to aid collaborative and multidisciplinary research. Through integration of gene expression datasets with context-specific and context-agnostic molecular interaction datasets, the workflow can be easily applied to different datasets as they are made available. Application to diverse SARS-CoV-2 datasets will increase our understanding of the mechanistic details of the infection at a cell type specific level, aid drug target discovery and help explain the variety of clinical manifestations of the infection.

## Introduction

By mid-May 2020 at least 4000 scientific preprints and publications were released relating to Severe Acute Respiratory Syndrome coronavirus 2 (SARS-CoV-2) and the disease it causes (COVID-19) (Kwon 2020). This fast uptake in research efforts is vital to decrease the health and economic impacts of this new pandemic. However, many questions remain unanswered regarding the molecular processes driving host responses to this coronavirus. One key challenge to utilisation of new findings is that published datasets are mostly unlinked to each other (due to parallel efforts by multiple research groups) and not always connected to community standard resources. An integrated and reusable method to interactively capture new data and connect it to existing data sources is needed. Such a comprehensive approach that can be run regularly when relevant new data is available, will increase and update our understanding of the mechanistic details of the SARS-CoV-2 infection. Further, it will aid drug target discovery by enabling identification of high confidence mediators through which the virus is affecting host cells (Barabási et al. 2011). Studying the effect of the virus at molecular level may explain the variety of clinical manifestations of the infection and the differences in susceptibility between different populations, and together with soon available human genomics data, could be used for identifying risk factors.

Upon entry of a virus into a human cell *via* surface receptors, viral RNA is released and translated into proteins (Oberfeld et al. 2020). In addition to their role in direct viral replication, these proteins are able to bind to human proteins creating a host-virus interface (Gordon et al. 2020). This interaction can lead to downstream signalling changes in the host cell, either as a result of viral hijacking or through a defined viral immune response by the host cell (Alto and Orth 2012). Ultimately, this signal flow results in intracellular gene transcription changes, cell-cell signalling and systemic host responses which drive the tug-of-war between the host and the virus (Fung et al. 2020). In order to understand and control this conflict, it is necessary to study each of these levels of host response in detail, including the intracellular response of the primarily infected cell.

Currently available data relating to intracellular SARS-CoV-2 infection includes human binding partners of viral proteins (Gordon et al. 2020) and transcriptomics datasets from infected cell lines/organoids (Blanco-Melo et al. 2020; Lamers et al. 2020), infected patients (Liao et al. 2020; Huang et al. 2020) and other infected animals (Pfaender et al. 2020; Blanco-Melo et al. 2020). Interdisciplinary and collaborative science can maximise the value of each of these datasets through data integration and comparison combined with application of different computational analysis approaches. One such computational analysis method is the utilisation of network approaches to model molecular interactions between the virus and human proteins as well as within and between human cells (Guven-Maiorov et al. 2017). Network approaches have already been applied to study SARS-CoV-2 pathogenesis and to predict drug repurposing candidates and master regulators based on proteins in proximity to human binding proteins (which physically associate with SARS-CoV-2 proteins) (Gysi et al. 2020; Messina et al. 2020; Zhou et al. 2020; Guzzi et al. 2020).

Here we present a systems biology workflow, to study the effect of viral infections on host cells. ViralLink reconstructs and analyses a causal molecular interaction network whose signal starts with the binding of an intracellular viral protein to a human protein, travels via multiple signalling pathways, and ends at the transcriptional regulation of altered genes. Subsequently, the workflow investigates the causal network using betweenness centrality measures, cluster analysis, functional overrepresentation analysis and network visualisation. Using currently available datasets from SARS-CoV-2 infected bronchial epithelial cells we demonstrate that this workflow can identify biologically relevant signalling pathways and predict key proteins for potential drug interventions. As the workflow is built in a modular, standardised and updateable fashion, it can be used easily in the future to analyse new SARS-CoV-2 related datasets (from human biopsy data, multiple tissues, etc.).

## Methods

### ViralLink workflow overview

The ViralLink workflow investigates the effect of viral infection within cells by generating and analysing context-specific networks of intracellular signalling and regulatory molecular interactions. These networks link the intracellular binding of viral and human proteins to the transcriptional response of the infected cell (Figure 1). The context-specificity of the analysis is obtained through the choice of input transcriptomics datasets - it could refer to strain of virus, type of infected cell, severity of infection, age of host or any other context of interest. By default, the workflow is set up to analyse the intracellular effects of SARS-CoV-2, requiring only transcriptomics counts data as input and thus encouraging and enabling rapid multidisciplinary research. However, the wide-ranging applicability and modularity of the workflow facilitates customisation of viral context, *a priori* interactions and analysis methods. ViralLink contains three primary stages: 1) collection and input of data; 2) reconstruction of the network; and 3) investigation of results using functional analysis, clustering, centrality measures and visualisation.

**Figure 1:**
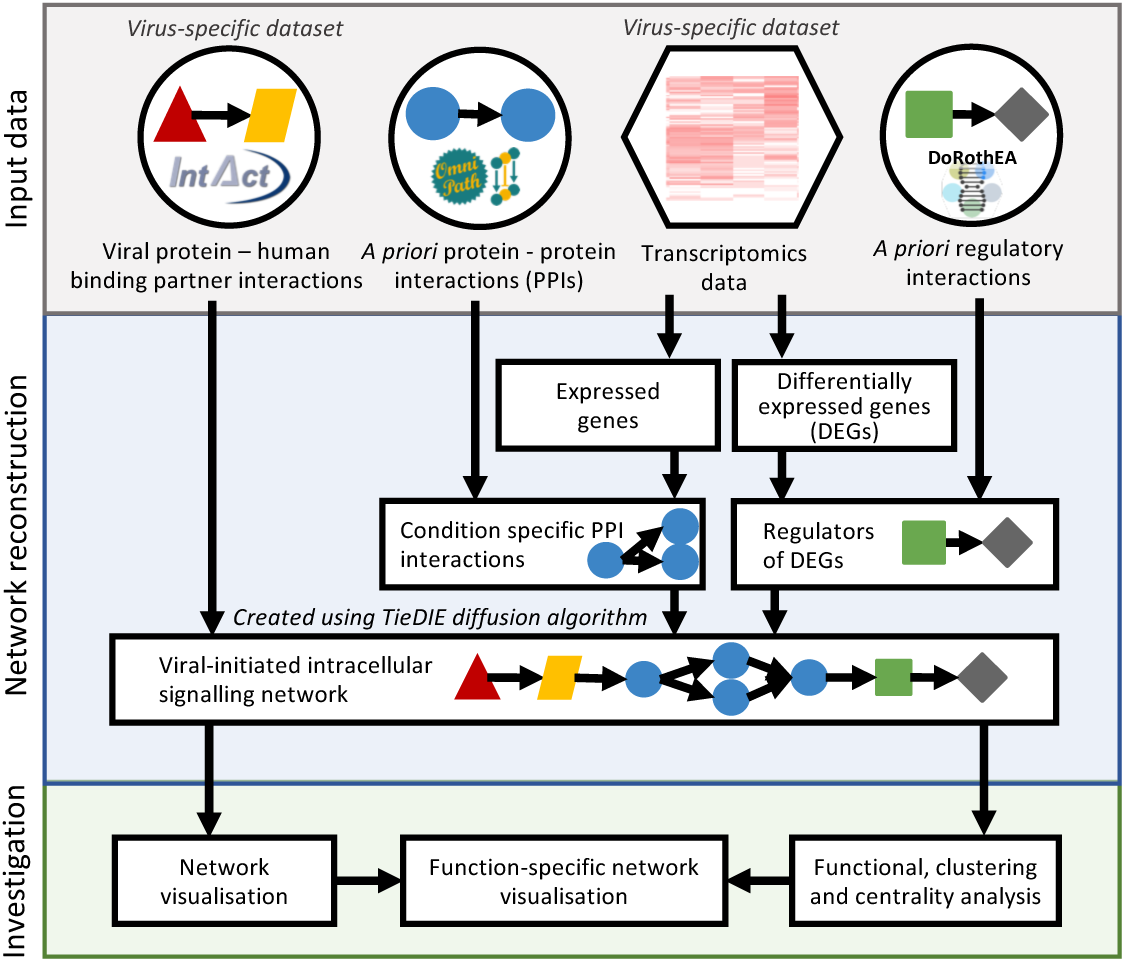
ViralLink workflow overview.

### Collection and input of data

Reconstruction of causal networks using ViralLink requires four separate input datasets (Figure 1): viral protein-human binding protein interactions, *a priori* human protein-protein interactions (PPIs), *a priori* human transcription factor (TF) - target gene (TG) interactions and an unnormalised counts matrix from a gene expression experiment. By default, all data except the transcriptomics counts are provided automatically. However alternative input files can be provided if desired.

The default workflow uses SARS-CoV-2 protein-human binding protein interactions obtained from an affinity-purification mass spectrometry study (Gordon et al. 2020) via Intact (Hermjakob et al. 2004; Orchard et al. 2014). This data was reformatted to contain one row per molecular interaction with 2 columns of UniProt IDs: SARS-CoV-2 proteins and human binding proteins. Alternative viral-human PPIs can be provided using the same data format. The workflow assumes all viral-human interactions have an inhibiting action on the human protein, unless a third column named “sign” is present in the input file containing “+” for activatory and “-” for inhibitory interactions. In addition, data is provided with the workflow containing the gene names corresponding to each of the SARS-CoV-2 proteins, to enable easy interpretation of the reconstructed networks.

For *a priori* human interactions, the workflow obtains and uses integrated collections of PPI and TF-TG interactions from OmniPath and DoRothEA, respectively (Türei et al. 2016; Garcia-Alonso et al. 2019). These interactions are obtained using the ‘OmniPathR’ R package (Türei et al. 2016; R Core Team 2013) to download and filter signed and directed interactions. For DoRothEA, only high and medium confidence level interactions are used (confidence scores A-C). In contrast to importing static input files, this script enables the use of up to date interaction data. Alternative interaction data can be used with the workflow provided it has the same format: specifically, it must contain source and target uniprot IDs in the columns ‘to’ and ‘from’ and if the transcriptomics data uses gene symbols, the interaction data must additionally contain gene symbols in the columns ‘source_genesymbol’ and ‘target_genesymbol’. Furthermore, the interactions must be directed and signed with the sign of the interaction given in the column ‘consensus_stimulation’ where the value ‘1’ represents a stimulation and anything else represents an inhibition.

The aforementioned *a priori* interactions are contextualised using transcriptomics data from any study of interest which compares viral infected to uninfected human cells or tissues. Correspondingly, the workflow requires unnormalised counts data from a transcriptomics experiment (containing Uniprot or gene symbols as IDs) and a corresponding mapping table which lists the sample IDs (from the headers of the counts table) in the ‘sample_name’ column and the ‘test’ or ‘control’ status of the sample in the ‘condition’ column. This mapping table is used to carry out differential expression of a test condition (e.g. infected) compared to a control condition (e.g. uninfected). An example expression dataset and mapping table are provided with the workflow.

To process the transcriptomics data, the workflow uses ‘DESeq2’ in R to normalise the counts and to carry out differential expression analysis (Love et al. 2014). Any genes passing the log2 fold change and adjusted p value cutoffs, based on the provided parameters (default 1 and 0.05, respectively), are classed as differentially expressed genes (DEGs). Following removal of all genes with count = 0, normalised log2 counts across all samples are fitted to a gaussian kernel (Beal 2017). All genes with expression values above mean minus three standard deviations are considered as expressed genes. Subsequently, context-specific human PPI and TF-TG interactions are generated by filtering only interactions where both interacting molecules are expressed.

File paths to all input datasets and associated parameters (such as desired log2 fold change cut off) are specified in the parameters text file which is read in by the workflow.

### Network reconstruction

The reconstructed causal network contains three layers of interactions, which are obtained, by default, from the three *a priori* interaction resources:

- Viral proteins interacting with human binding partners: from the SARS-CoV-2 collection in the IntAct database (Hermjakob et al. 2004; Orchard et al. 2014)
- Intermediary signalling protein interactions: from protein-protein interactions (PPIs) of the OmniPath collection (Türei et al. 2016)
- Transcription factors (TFs) regulating differentially expressed genes: from a transcriptomics dataset of interest and the DoRothEA collection (Garcia-Alonso et al. 2019)

A list of all TFs targeting the differentially expressed genes are obtained from the context-specific TF-TG interactions. The human binding proteins of viral proteins are connected to the listed TFs through the context-specific human PPIs using a network diffusion approach called Tied Diffusion Through Interacting Events (TieDIE) (Paull et al. 2013). As inputs for the TieDIE tool, the following information is used: (1) The signed, directed and expression based filtered PPIs is used as the input network. (2) Human proteins which are interacting partners of the viral proteins are used as the start nodes. The number of viral proteins bound to each of the human proteins are assigned as the weights of the start nodes. (3) The TFs of the DEGs in the dataset are used as the stop nodes. The weights for each of the TFs in the set of stop nodes were calculated using the following formula (Equation 1) which considers both the log2 fold change of the DEGs as well as the sign (i.e stimulatory or inhibitory) of the relationship between the TF and the DEG.

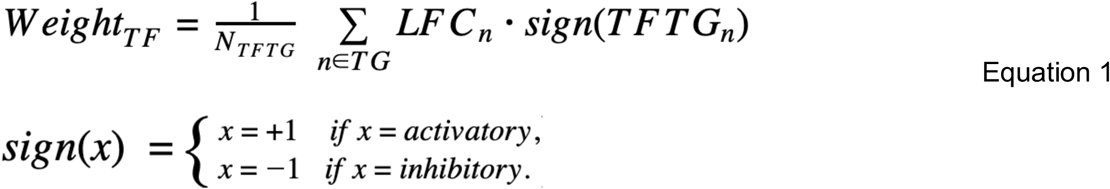

After running TieDIE, a custom R script is used to collate all the data into a final viral-initiated intracellular signalling network (causal network), outputting an edge table representation of the network, with a node table containing additional node annotations. Starting with the interactions output from TieDIE, viral protein-human binding protein interactions are added for each of the present human binding proteins. Similarly, TF-TG interactions (where the TG is a DEG) are added for each of the present TFs, creating a full network with three interaction types: SARS-CoV-2 protein-human binding protein, PPI and TF-DEG. All nodes of the network are added to a node table with annotations including heat values (output from TieDIE), Entrez IDs (obtained in R using the ‘org.Hs.eg.db’ package), gene symbols (obtained from UniProt (UniProt Consortium 2019)) and log2 fold change values from the differential expression analysis.

### Network investigation

Following reconstruction of the causal network, ViralLink provides functionality to investigate the results using functional analysis, clustering, centrality measures and visualisation.

#### Centrality measures

To identify key molecules in the reconstructed network ViralLink uses a betweenness centrality measure - calculating the global importance of a node (in this case a protein) based on the number of shortest paths which pass through them when connecting all node pairs in the network (Koschützki and Schreiber 2008). Nodes with high betweenness centrality play a key role in transduction of signals through the network, and here represent proteins with biological importance in the cellular response to viral infection. Betweenness centrality is calculated for each node in the causal network using the R package ‘igraph’ and output as an annotation in the node table (Csárdi and Nepusz 2006). Alternative centrality measures are available using the ‘igraph’ package and can be integrated into the workflow by the user if required.

#### Cluster analysis

Clustering algorithms are commonly used in network biology to investigate the complex structure of molecular interaction networks by extracting groups of densely connected molecules (Bader and Hogue 2003; Brohée et al. 2008). Depending on the number of molecules included, a cluster can represent a molecular complex or a group of molecules which function closely with each other. Cluster analysis can identify subsets of a large network with specific functions and indicate molecules that may have functional redundancy with each other - potentially having implications for drug targeting. ViralLink employs the MCODE clustering method to identify groups of densely connected nodes in PPI networks (Bader and Hogue 2003). To carry out this analysis, ViralLink requires a local version of the Cytoscape software to be open (Shannon et al. 2003; Su et al. 2014), which is controlled programmatically using the R package ‘RCy3’ with the Cytoscape ‘MCODE’ app (v1.6.1) (Gustavsen et al. 2019). MCODE is run using default parameters: degree cut off =2, haircut=TRUE, node score cut off=0.2, k-core=2, max depth=100. This analysis outputs the data as node annotations in the node table, which are used for the functional analysis and visualisation steps of the workflow. If Cytoscape is not running, this step of the workflow will be skipped.

#### Functional analysis

To further investigate important cellular functions and signalling pathways directly affected by the virus of interest, ViralLink carries out functional overrepresentation analysis on different parts of the causal network:

1. The DEGs of the network
2. The upstream human proteins (including human binding proteins, intermediary signalling proteins and TFs)
3. Identified clusters (only those with ≥ 15 nodes are investigated)

Functional overrepresentation analysis is carried out in R using packages ‘ClusterProfiler’ (for Gene Ontology annotations (Ashburner et al. 2000)) and ‘ReactomePA’ (for Reactome annotations (Yu et al. 2012; Yu and He 2016; Fabregat et al. 2018). For analysis of the upstream human signalling proteins and analysis of clusters, all proteins in the context-specific human PPI interactions are used as the background. For analysis of the DEGs, all target genes in the context-specific human TF-TG interactions are used as the background. For Gene Ontology (Biological Process) analysis (except when running the compareCluster command), the ‘simplify’ command is used (cutoff=0.1, select_fun=min) to remove redundant functions. All functions with q val ≤ 0.05 are considered significantly overrepresented.

An additional R script is provided alongside the workflow which creates subnetworks of the causal network based on functions of interest. These function-specific subnetworks highlight how specific signalling pathways in the infected cell reach (and subsequently affect) specific functions of the DEGs. For example, the subnetwork could be created to show how viral proteins can affect different host toll-like receptor pathways, and how these pathways can ultimately affect DEGs associated with interleukins. In this network the DEG nodes would be replaced with nodes representing the interleukin functions (which must be overrepresented based on the functional analysis). This script requires the output files from the functional analysis, the node and edge tables of the causal network and a file of all Uniprot IDs associated with all Reactome functions (which is provided with ViralLink, following download from the Reactome website in April 2020). In addition, the script requires a list of overrepresented DEG functions (Reactome) and a list of upstream signalling functions (Reactome) to visualise. The script outputs an edge table, a node table and a Cytoscape file (if Cytoscape is open locally at the time of running the script).

#### Visualisation

Data visualisation is often an important part of biological network interpretation, providing new insights into the data and visually conveying analysis results (Pavlopoulos et al. 2008). As such, ViralLink has the capability to import reconstructed networks into the open-source Cytoscape network visualisation software (Shannon et al. 2003; Su et al. 2014). This functionality requires that the user has Cytoscape installed and open locally. Specifically, the workflow employs the ‘RCy3’ R package to interact with Cytoscape programmatically, importing the node and edge tables to create network visualisations and saving the data as a ‘.cys’ file. The causal network, the network clusters (where containing ≥ 15 nodes) and the function-specific networks are visualised in this way. If calculated previously, the causal network nodes are coloured based on their betweenness centrality, however further style and layout customisation must be carried out by the user directly based on the data.

### Implementation

The workflow consists of modular R and Python scripts which can be run separately or through the provided Python wrapper script. If running for the study of SARS-CoV-2, the only required input files are related to the transcriptomics data of interest: a raw counts table (using gene symbols or UniProt protein IDs) and a two-column metadata table specifying test and control sample IDs. One further script is provided to generate function-specific networks. This script is not included in the wrapper because it requires the user to specify functions of interest from the output of the functional analysis. To run everything, it is necessary that the user has R, Python3 and Cytoscape installed. The only file the user needs to edit is the parameters text file where input file paths and parameters are specified. All scripts, default input files and details of how to run the scripts are freely accessible on GitHub (https://github.com/korcsmarosgroup/ViralLink).

### Use case

To demonstrate the application of this workflow for the study of SARS-CoV-2, we applied it to a published transcriptomics dataset. We downloaded raw counts tables from a transcriptomics study of SARS-CoV-2 infected (MOI 2, 24 hour incubation) NHBE cells (Normal Human Bronchial/tracheal Epithelial cell line) with uninfected controls (Blanco-Melo et al. 2020) via Gene Expression Omnibus (accession GSE147507) (Edgar et al. 2002; Barrett et al. 2013). OmniPath and DoRothEA (v2, A-C) were downloaded on 15/04/2020. Any genes with log2 fold change ≥ |0.5| and adjusted p value ≤ 0.05 were classed as differentially expressed. All networks were visualised in Cytoscape (v3.7.2).

## Results

### Use case: SARS-CoV-2 infection of lung cells

To demonstrate the application of this workflow for the study of SARS-CoV-2, we created intracellular signalling networks of NHBE cells (from Normal Human Bronchial/tracheal Epithelial cell lines) upon infection with SARS-CoV-2 based on data published by Blanco Melo *et al*. (Blanco-Melo et al. 2020) and viral-human binding protein interactions published by Gordon *et al*. (Gordon et al. 2020). The resulting causal network contains 804 nodes (molecules) and 5423 interactions (Figure 2A, Supplementary Tables 1-2, Supplementary File 1). The 10 most central proteins of the reconstructed causal network (based on betweenness centrality) are involved in a wide range of cellular functions (Figure 2B). Taken together these proteins highlight the propensity for SARS-CoV-2 to affect cell proliferation, apoptosis, cell adhesion, exocytosis and proinflammatory immune responses. These functions are influenced through multiple cellular pathways, most notably MAPK/ERK and PI3K/AKT signalling pathways.

**Figure 2.**
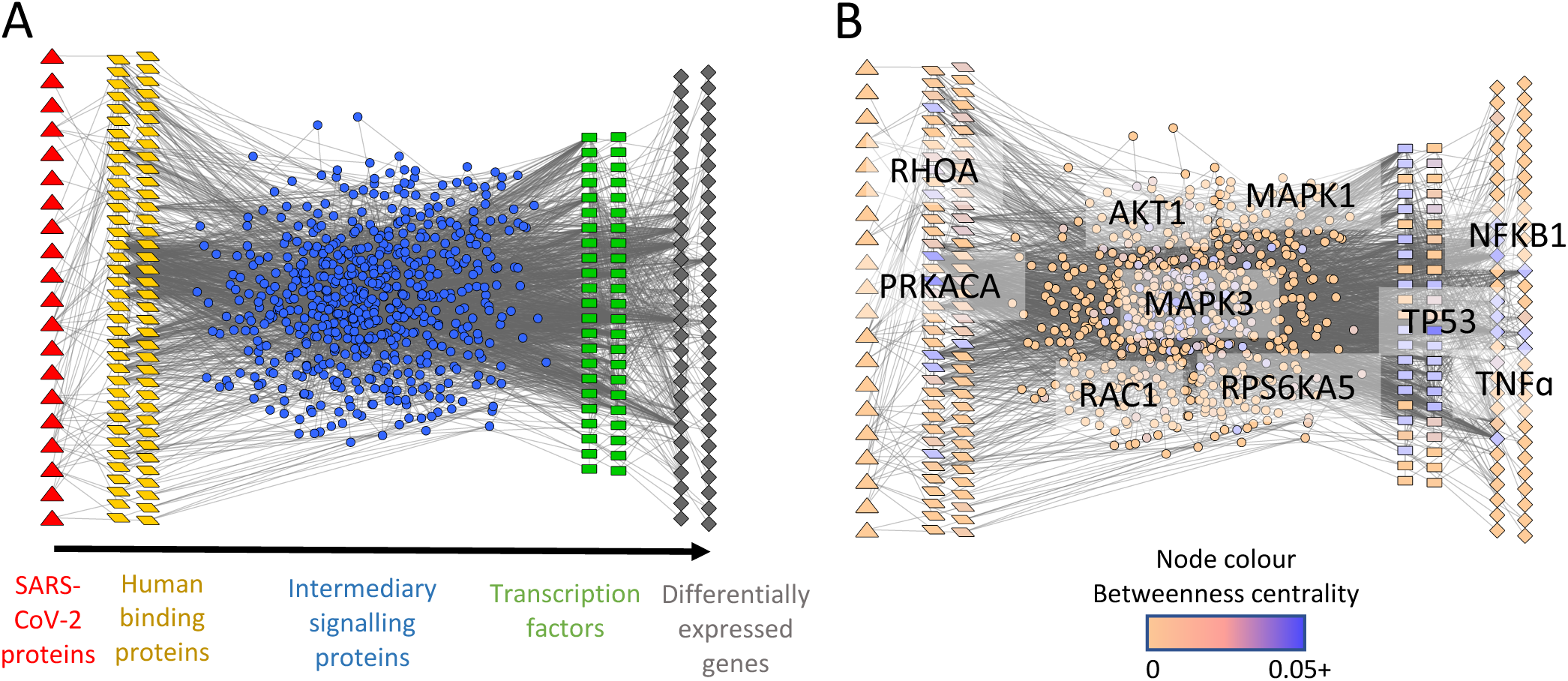
Causal network of SARS-CoV-2-infected NHBE cells. **A)** Signalling flows from left to right: SARS-CoV-2 proteins/protein fragments (red triangles), human binding proteins (yellow parallelograms), intermediary signalling proteins (blue circles), transcription factors (green rectangles) and differentially expressed genes (grey rhombuses). Where a human protein/gene is acting in multiple layers of the network, it is only visualised once based on the following priority: DEGs, binding proteins, TFs, signalling proteins. **B)** Results of betweenness centrality analysis, which measures the global importance of nodes (molecules) in the network. Nodes coloured based on their betweenness centrality parameter, with the gene names of the 10 highest scoring (most central) nodes overlaid. DEGs have log2 fold change ≥ |0.5| and adjusted p value ≤ 0.05.

Functional overrepresentation analysis of the causal network identified an enrichment of interleukin and interferon related functions among the network DEGs, in line with previously published findings (Supplementary Figure 1, Supplementary File 2) (Zhang et al. 2020; Chua et al. 2020; Huang et al. 2020). Overrepresented functions and pathways of the upstream signalling proteins (human binding proteins, intermediary signalling proteins and TFs) included innate immunity-related functions, platelet signaling, PI3K/AKT signalling, MAPK activation, estrogen receptor-mediated signalling, senescence and a number of growth factor receptor-associated functions (such as VEGF signalling, receptor tyrosine kinases, stem cell growth factor signalling (SCF-KIT) and neurotrophin receptor signaling). Therefore, we show that this analysis highlights additional pathways through which SARS-CoV-2 could be affecting the lung epithelial cells, which cannot be identified by looking at the transcriptomic results in isolation.

Based on functional overrepresentation analysis, we created a function-specific network by sub setting the causal network. This visualisation was used to further explore the mechanisms of how specific signalling pathways are affecting the DEGs (Supplementary Figure 2A, Supplementary File 3). Specifically, we generated an innate-immunity associated subnetwork containing all upstream human signalling proteins associated with Reactome functions cytokine signalling in immune system, signaling by interleukins and MyD88-independent TLR4 cascade and all overrepresented functions of the DEGs (in place of the DEG nodes). These pathways contain 9/10 of the top betweenness centrality nodes (all except RHOA), evidencing the centrality and importance of the innate immune response to viral infection. Inspecting the TF layer of this immune subnetwork, we find a number of key TFs including STAT proteins (3 and 4), IRF proteins (1 and 5) and NFKB-related proteins (NFKB1, NFKBIA).

Finally, we evidenced the application of MCODE clustering analysis to using the reconstructed SARS-CoV-2-infected NHBE cell causal network. We identified four clusters containing 15 or more nodes, making up 19% of the network (154/804) (Supplementary Figure 2B, Supplementary Table 2, Supplementary File 1). Assuringly, 9/10 of the top betweenness centrality nodes were included in these four clusters, further confirming the high connectivity and importance of these nodes in the causal network. Functional overrepresentation analysis of the cluster nodes highlighted a functional similarity between all four of the clusters (Supplementary Figure C-D, Supplementary File 2). Likely this is due to the high number of inter-cluster molecular interactions and because of the functional similarities between the top central nodes.

Collectively, we show that our systems biology workflow, ViralLink, reconstructs a functionally relevant intracellular signalling network affected by SARS-CoV-2 infection. Investigation of the networks through functional analysis, centrality measures and cluster analysis, combined with network visualisations, enables detailed study of the key proteins and pathways involved in signal transduction.

## Discussion

Infection by SARS-CoV-2 can cause a complex and systemic response by the human body. As such, a better mechanistic understanding of the effects of SARS-CoV-2 will aid identification of effective drug treatments and help to explain the differences in susceptibilities across different populations (Kirby 2020). This understanding can be gained using cross-disciplinary approaches which combine ‘omics data generation, computational systems biology and validatory web lab experiments (Korcsmaros et al. 2017). Here we present a computational workflow that can be used to model the cellular response to infection by integrating knowledge of human binding proteins of viral proteins with the transcriptional response of a cell/cell type. Whilst set up primarily to run analyses based on SARS-CoV-2, ViralLink can be applied to any viral infection, provided data is available describing possible interactions between the viral proteins and human proteins.

ViralLink builds on our previously published resource MicrobioLink, which reconstructs networks representing the effect of extracellular and intracellular microbial proteins on cellular processes (Andrighetti et al. 2020). Differing from MicrobioLink, ViralLink inputs a predetermined list of viral-host PPIs and focuses only on pathways ending in transcriptional regulation: thereby reducing the complexity of the workflow (for accessibility and speed purposes) and increasing its predictive confidence. Furthermore, ViralLink extends the functionality of MicrobioLink with more advanced network analysis (functional enrichment, clustering and centrality measures) and visualisation options.

By exploiting previously collated and comprehensive collections of molecular interactions (Türei et al. 2016; Garcia-Alonso et al. 2019), ViralLink predicts how signal flows from the initial interaction with a viral protein or protein fragment to the ultimate transcriptional changes induced by the virus. Through mapping the direct intracellular effect of viral infection (using a network approach), this workflow enables further investigation into specific signalling pathways and transcription factors which play a key role in signal transduction. Signalling pathways are primarily regulated through post-translational modifications and thus would not be identified using transcriptomics datasets (Antebi et al. 2017). In addition, the resulting intracellular networks allow identification of differentially regulated genes that are affected as a direct result of viral recognition by protein-protein signalling pathways, rather than by secondary signals such as elevated cytokine levels. This permits a more focused analysis of possible drug targets and adds to the understanding of viral pathomechanisms. Functional analysis and visualisation methods included in the workflow are vital for interpretation of the generated intracellular networks, enabling detailed investigation of key proteins and signalling pathways.

Due to the modularity of the workflow, it can be easily adjusted or extended - different diffusion and propagation algorithms, such as HotNet2 (Leiserson et al. 2015; Cowen et al. 2017), could be implemented as required. The implemented diffusion tool, TieDIE, adds mechanistic value by accounting for local causality (e.g. sign) but, on the other hand, has a reduced possible set of input *a priori* interactions. If desired, a diffusion tool which does not need signed *a priori* interactions can be implemented to increase the input dataset size. Alternatively, a different method, such as an integer linear programming approach which identifies paths based on an optimisation problem (as implemented in CARNIVAL), could be used for network reconstruction (Liu et al. 2019). In addition, integration of CARNIVAL could extend the workflow to permit network reconstruction without supplying upstream perturbations (in this case the viral-host protein interactions). Whilst not currently integrated due to data availability issues, the addition of phosphoproteomics data to the pathway propagation methods could improve the prediction of active pathways (Dugourd et al. 2020) Alternatively, methods to predict protein activity based on transcriptional signatures, such as VIPER and PROGENy (Alvarez et al. 2016; Schubert et al. 2018) could be added to the workflow in addition to network diffusion methods to increase the confidence of pathway predictions. Finally, extension of the network to include additional regulatory molecule types (e.g. miRNAs) or to study non-human hosts, could uncover further mechanisms by which SARS-CoV-2 can affect host cells.

Accessible through GitHub, the workflow requires R and Python3 to be installed (and Cytoscape for clustering and visualisation), however only a limited programming ability is required to run the code. All code is wrapped into a Python script with a separate file where all input file paths and parameters are specified. At a minimum, only two user specified input files are required: a raw counts table from a transcriptomics study (using gene symbols or UniProt protein IDs) and a two-column metadata table specifying test and control sample IDs. All other files are provided or acquired directly within the workflow - but can be changed by the user if required. However, one limitation of the current workflow is that creation of Cytoscape visualisations and clustering analysis require the user to install and open the Cytoscape app. If this is not possible, for example because the scripts are not being run on a machine with a graphical interface, these steps are skipped. Furthermore, only basic visualisation is possible programmatically, due to challenges applying one visualisation strategy to all possible output networks, especially with regard to the function-based networks.

In addition to accessibility through a default emphasis on SARS-CoV-2, a key strength of this workflow is the ability to use different input datasets: including different *a priori* molecular interactions, viral-human binding protein interactions and expressed/differentially expressed gene lists. This allows extensive customisation and permits rapid implementation to the most cutting-edge data soon after publication. Running the workflow across different transcriptomics datasets will allow comparison of intracellular viral responses between different cell types, different species and across different conditions (such as severe vs asymptomatic infection). For example, application of the workflow to transcriptomics data from specific immune cell-types, such as macrophages, will likely uncover different host affected signalling pathways and key TFs based on the infected cell-type. This, in turn, could increase our understanding of the role of different immune populations in fighting the infection. In addition, the workflow can be run on data from other SARS-CoV-2 strains when and if they emerge, thereby aiding comparisons of mechanisms of action between the strains.

To evidence the use of this workflow, we applied it to study the effect of SARS-CoV-2 infection in lung epithelial (NHBE) cells using transcriptomics data published by Blanco-Melo *et al*. (Blanco-Melo et al. 2020). In the resulting causal network, DEGs directly affected by SARS-CoV-2 initiated signalling are associated with functions that are known responses to SARS-CoV-2 and other viral infections (Cao 2020; Shi et al. 2020; Sallard et al. 2020; Arvanitakis et al. 1998). Upstream of these affected genes we identified a number of potentially important signalling pathways relating to classical viral-immune responses, cell survival and cytoskeletal rearrangements and cell adhesion. Previous investigation of the first SARS coronavirus (SARS-CoV) identified an inhibition of cell proliferation and an increase in apoptosis regulated to PI3K/AKT signalling (Mizutani et al. 2006; Tsoi et al. 2014). Our network of SARS-CoV-2-initiated intracellular signalling suggests that the PI3K/AKT signalling and the AKT1 protein itself are key mediators of SARS-CoV-2 initiated signal transduction and that apoptosis and cell proliferation pathways are affected by SARS-CoV-2, thus highlighting similarities between the two viruses. However, further experimentation and/or data curation is required to confirm the direction of change of specific pathways (up- or downregulated) based on the results of the presented workflow. Together our results indicate that SARS-CoV-2 can affect NHBE cells through a variety of signalling pathways which have been previously associated with similar viruses, including growth factor signalling, MAPK/ERK signalling and PI3K/AKT signalling. Furthermore, centrality measures and cluster analysis identified proteins which likely play a key role in transduction of these signals, and could be good targets for drug treatments.

Several other network reconstruction methods exist which could be and have been applied to study SARS-CoV-2 infections. For example Messina *et al*. and Gysi *et al*. (Messina et al. 2020; Gysi et al. 2020) use diffusion algorithms and other similar methods to investigate proteins in close proximity to human binding proteins based on PPI interactions and gene co-expression networks. Our workflow builds on these approaches by linking viral proteins to DEGs. Through this method we can observe which signalling pathways mediate the effect of the virus on cellular transcription levels, creating a systems level view of cellular changes as a result of the virus. Using the functional analysis methods and network visualisation capabilities of the workflow, it is possible to predict which viral proteins and host signalling pathways can affect specific cellular functions, enabling more focused identification of drug targets. In addition to protein mediators, this method describes TFs which are involved in the cellular response and identifies which DEGs can be affected as a direct result of viral proteins hijacking host signalling and which are affected through a different mechanism. In addition to the presented workflow, at least one other method has been used to reconstruct SARS-CoV-2-initiated intracellular signalling networks (Ding et al. 2020) corroborating the benefits of such analysis methods. Differing from the here presented approach, this work uses an extended version of the Signaling Dynamic Regulatory Events Miner method to reconstruct the networks, resulting in a more mathematically complex but computationally heavy analysis (Gitter et al. 2013). Furthermore, the workflow by Ding *et al*. is a less reusable and accessible workflow because it was designed for a specific analysis.

In conclusion, ViralLink is an easily accessible, reproducible and scalable systems biology workflow to reconstruct and analyse molecular interaction networks representing the effect of the viruses on intracellular signalling. We believe it is the first available integrative workflow for analysing the downstream effects of viral proteins using viral host interactions and host response data. Application of this workflow to study COVID-19 based on a wide variety of conditions and datasets will uncover mechanistic details about SARS-CoV-2 infection of different cell types, providing valuable predictions for wet-lab and clinical validation.

## Acknowledgements

Many thanks to members of the Saez-Rodriguez group and to the COVID-19 Disease Map Community for their ideas and support. In particular we thank Julio Saez-Rodriguez, Alberto Valdeolivas and Aurelien Dugourd for their advice and discussions and Nigel Fosker for help with the manuscript. This research was supported in part by the NBI Computing infrastructure for Science (CiS) group through the provision of a High-Performance Computing (HPC) Cluster.

## Funding

A.T., L.G., M.O., M.P. and M.M. are supported by the BBSRC Norwich Research Park Biosciences Doctoral Training Partnership (grant numbers BB/M011216/1 and BB/S50743X/1). The work of T.K., D.M., and I.H. was supported by the Earlham Institute (Norwich, UK) in partnership with the Quadram Institute (Norwich, UK) and strategically supported by the UKRI Biotechnological and Biosciences Research Council (BBSRC) UK grants (BB/J004529/1, BB/P016774/1, and BB/CSP17270/1). T.K. and D.M. were also funded by a BBSRC ISP grant for Gut Microbes and Health BB/R012490/1 and its constituent project(s), BBS/E/F/000PR10353 and BBS/E/F/000PR10355. P.S. was supported by funding from the European Research Council (ERC) under the European Union’s Horizon 2020 research and innovation programme (grant agreement no. 694679).

## Figure and Tables

**Supplementary Figure 1.**
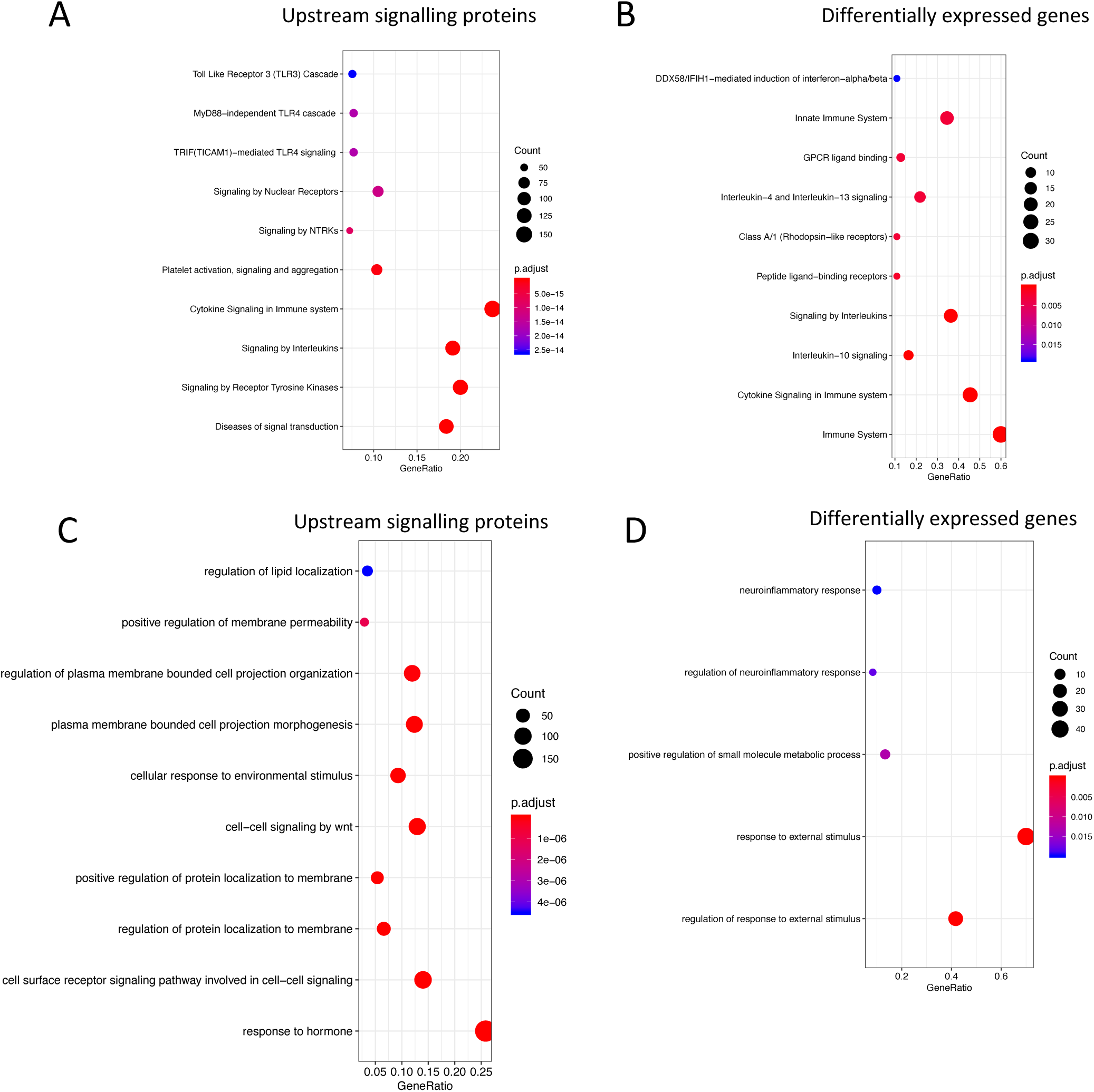
Overrepresented Reactome functions (A, B) and Gene Ontology Biological Processes (C, D) of the causal network of SARS-CoV-2 infected NHBE cells. **A)** Top 10 overrepresented Reactome functions of upstream signalling proteins (including human binding proteins, intermediary signalling proteins and TFs) **B)** Top 10 overrepresented Reactome functions of network DEGs C) Top 10 overrepresented GO-BP functions of upstream signalling proteins (including human binding proteins, intermediary signalling proteins and TFs) D) All overrepresented GO-BP functions of network DEGs (q value ≤ 0.05). DEGs have log2 fold change ≥ |0.5| and adjusted p value ≤ 0.05.

**Supplementary Figure 2:**
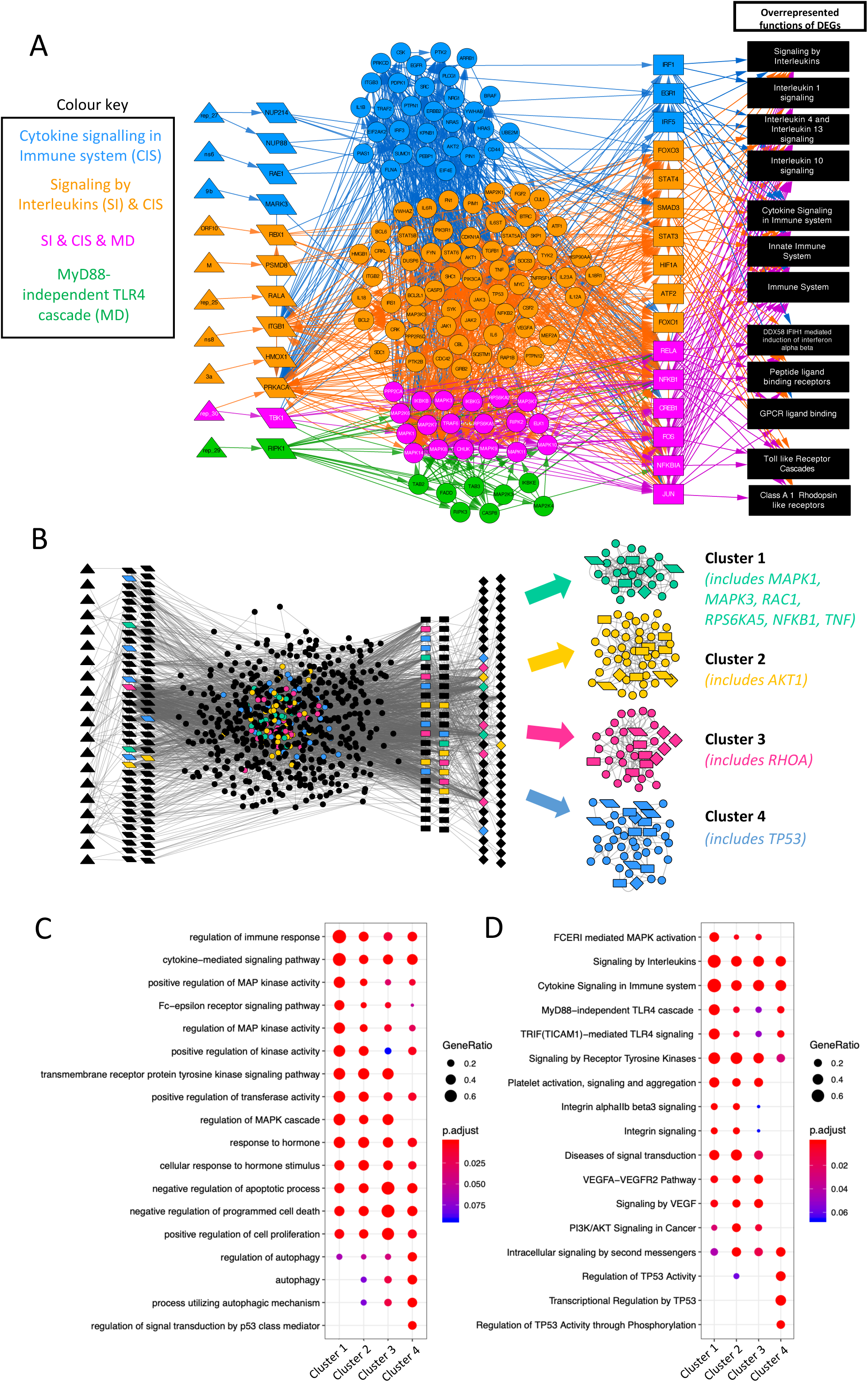
Function-specific network SARS-CoV-2-infected NHBE cells and cluster analysis on SARS-CoV-2-infected NHBE causal network. **A)** Function-specific subnetwork containing upstream signalling proteins related to the top overrepresented (q value ≤ 0.05) innate immunity-related Reactome functions (cytokine signalling in immune system, signaling by interleukins and MyD88-independent TLR4 cascade) and all overrepresented functions of the DEGs (in place of the DEG nodes). Layers of the network and node shapes same as in Figure 2. DEGs = differentially expressed genes. DEGs have log2 fold change ≥ |0.5| and adjusted p value ≤ 0.05. See Supplementary File 3. B) Cluster analysis results where clusters have ≥ 15 nodes. Position of clustered proteins shown within the causal network and to the right as isolated clusters. Nodes coloured by their cluster membership (black=unclustered, green=cluster 1, yellow=cluster 2, pink=cluster 3, blue=cluster 4). Presence of top 10 betweenness centrality nodes in the clusters is indicated to the right of the clusters. B) Gene Ontology (GO) overrepresentation analysis of the clusters. Top five GO terms (by adjusted p value) displayed for each cluster. **C)** Reactome overrepresentation analysis of the clusters. Top five Reactome terms (by adjusted p value) displayed for each cluster. See Supplementary Table 2 and Supplementary File 2.

**Supplementary Table 1: Causal network of SARS-CoV-2-infected NHBE cell**.

**Supplementary Table 2: Node annotations for causal network of SARS-CoV-2-infected NHBE cell**. Includes betweenness centrality measures and clusters identified by MCODE. MCODE clusters 1,3,4 and 5 correspond to the clusters in the manuscript labelled 1,2,3 and 4 respectively. Clusters 2 and 6 were excluded due to size.

**Supplementary File 1: Causal network of SARS-CoV-2-infected NHBE cell, Cytoscape file**.

**Supplementary File 2: Functional overrepresentation results**. Reactome and Gene Ontology Biological Processes (q value <= 0.05) for differentially expressed genes (DEGs), protein-protein (PPI) interaction nodes (human binding proteins, signalling proteins and transcription factors) and the clusters of the causal network of SARS-CoV-2-infected NHBE cell.

**Supplementary File 3: Function-specific network of SARS-CoV-2-infected NHBE cells, Cytoscape file**.

## Notes

### Competing Interest Statement

The authors have declared no competing interest.

### Summary of Updates

Updated author affiliations

